# Temperature dependent sex-biased gene expression in the gametophytes of the kelp *Saccharina latissima*

**DOI:** 10.1101/750455

**Authors:** Cátia Monteiro, Sandra Heinrich, Inka Bartsch, Klaus Ulrich Valentin, Erwan Corre, Jonas Collén, Lars Harms, Gernot Glöckner, Kai Bischof

## Abstract

*Saccharina latissima* is an economically and ecologically relevant kelp species in Europe and North America. In kelps, the sexuality is expressed during the haploid life stage and the microscopic gametophytes exhibit significant sexual dimorphism. To understand the sex-dependent impact of temperature on the gametophyte stage, we analyzed for the first time, gene expression profiles of male and female gametophytes at three different temperatures (4°C, 12°C and 20°C) characteristic for the species distribution range by using RNA-sequencing. We identified several differentially expressed genes between sexes; while female biased genes were enriched in general metabolism and energy production, male biased genes function within cell cycle and signaling. In our study, temperature modulated sex-biased gene expression, with only a small percentage of differentially expressed genes consistently male (7%) or female-biased (12%) at the three temperatures. Female gametophytes responded stronger to higher temperatures than males, suggesting that males are more heat tolerant. Differences between *S. latissima* and other brown algal gender-dependent gene expression might mirror the different evolutionary and ecological contexts. Genomic information on kelp gametophyte is still scarce and thus this study adds to our knowledge on sex differences in abiotic stress responses in macroalgae at the transcriptomic level.

**Highlight:** The transcriptomic basis for sexual dimorphism and associated metabolic needs are described for the kelp *Saccharina latissima*. Temperature modulates sex-biased gene expression resulting in a stronger stress response in females.

## Introduction

Kelps (order Laminariales, Phaeophyceae) feature a haplo-diplontic life cycle in which microscopic haploid gametophytes alternate with macroscopic diploid sporophytes. The two stages are very distinct: sporophytes are structured in holdfast, stipe and blade featuring different tissues (cortex and medulla) and may reach up to several meters in length, while gametophytes are composed of one to few cells (Hurd *et al.*, 2014b). These morphological differences between the life stages entail different susceptibility to abiotic stressors (Coelho *et al.*, 2000), namely high irradiance, ultraviolet radiation and temperature (e.g. Bolton and Lüning, 1982; Fredersdorf *et al.*, 2009; Gao *et al.*, 2019). Therefore, any prediction of future impacts of global warming on kelps must not only consider the responses of the sporophytes but also of the other life-history stages (Assis *et al.*, 2017; Leal *et al.*, 2018; Roleda, 2016).

Reproduction in kelps is tightly connected to environmental cues. Gametophytes may enter gametogenesis under the specific combination of nutrients, light and temperature or remain vegetative if conditions are not favorable (Bartsch, 2018; Martins *et al.*, 2017). In the latter case, vegetative gametophytes are considered to be dormant and may act as a “seed bank” allowing for the recovery of the population after a disturbance, e.g. storms or heat-waves (Barradas *et al.*, 2011; Edwards, 2000). Moreover, vegetative gametophytes might be more resistant than macroscopic sporophytes to heat stress during summer, especially at the rear edge of the distribution. Lee and Brinkhuis (1986) suggested that the gametophyte and/or early life stages of *Saccharina latissima* “oversummer” at the southern distribution limit in USA, as sporophytes are not visible during the hot summer months but are already found in early autumn. Hsiao and Druehl (1973) reported gametophyte survival and gamete production of *S. latissima* throughout the year, however absence of macroscopic sporophytes during later spring and summer. Given the challenges of studying the microscopic stages in the field, the relevance of the vegetative gametophyte pool for population dynamics is not well understood (Schiel and Foster, 2006). Nevertheless, manipulation of life-cycle transitions is a common practice in aquaculture to allow for year-round production of sporophytes (Charrier *et al.*, 2017). Control of gametogenesis starts by separating male and female gametophytes based on their sexual dimorphism. After spore release and germination, sexual dimorphism soon becomes visible: while female gametophytes are constituted of a few bigger rounder cells that tend to grow in diameter, male cells are much smaller and increase faster in cell numbers (Lüning and Neushul, 1978). Under non-fertilizing conditions, male and female gametophytes form vegetative filaments consisting of several cells that are able to continuously grow over decades (e.g.Martins *et al.*, 2019) and may form mm-sized clumps (Bartsch, 2018). The significance of sex to differences in performance, behavior and ecology are also currently understudied in brown algae (Luthringer *et al.*, 2014). Sexual dimorphism is to a large extent mediated by differential gene expression between sexes in a wide variety of organisms (Ellegren and Parsch, 2007). Up to the present, only a few gametophyte gene expression datasets are available for brown algae: *Ectocarpus spp.* (order Ectocarpales) (Lipinska *et al.*, 2015), *Undaria pinnatifida* (Shan *et al.*, 2015), *Saccharina japonica* (order Laminariales) (Ye *et al.*, 2015), *S. latissima* (GA Pearson *et al*. 2019, unpublished data). Transcriptomic differences between sexes in Phaeophyceae have been seldom explored, except for reproductive tissues in *Fucus vesiculosus* (Martins *et al.*, 2013), *S. japonica* gametophytes (Bi and Zhou, 2014) *Ectocarpus spp.* gametophytes and gametes (Lipinska *et al.*, 2015; Lipinska *et al.*, 2013) and in *S. latissima* gametophytes in response to gametogenesis induction and light quality changes (GA Pearson *et al*. 2019, unpublished data). Furthermore, so far, the influence of temperature has not been addressed. In the present study we provide the first data on transcriptomic responses of male and female gametophytes of the sugar kelp, *S. latissima*, to temperature. *Saccharina latissima* is a relevant foundation species in temperate to Arctic coasts (Steneck *et al.*, 2002; Teagle *et al.*, 2017) and it is the selected species in several aquaculture facilities in Europe and North America (Broch *et al.*, 2018; Forbord *et al.*, 2012; Kim *et al.*, 2015).

Our experiment targeted gametophytes of *S. latissima* originating from Helgoland (German Bight, North Sea) representing its center of distribution in respect to latitude (Araújo *et al.*, 2016). Despite this, summer temperatures observed at this site are high (up to 20°C) (Breitbach *et al.*, 2016) and close to the upper thermal tolerance limit of the species (Bolton and Lüning, 1982). Moreover, a decline in *S. latissima* relative presence and abundance has been described in Helgoland recently and were paralleled by increases in temperature and water clarity (Pehlke and Bartsch, 2008).

To our knowledge, this is the first study targeting brown algae gametophyte responses to temperature at the gene expression level. Hence, the present dataset will largely contribute to the understanding of the genomic basis of sex-specific differences in response to the environment of brown algae.

## Material and Methods

### Algal material and experimental design

Gametophytes from uniparental stock cultures of *Saccharina latissima* (formerly *Laminaria saccharin*a L. (Lamour) (Lane *et al.*, 2006) (AWI culture number 3094 (males) and 3096 (females)) from Helgoland, Germany were cultivated in glass flasks under red light (16hL:8hD) at 12°C (±1°C) in an environmentally controlled room. Sterile seawater enriched with an adapted Provasoli medium without the addition of iron was replaced weekly. These conditions prevented the onset of gametogenesis (Lewis *et al.*, 2013; Lüning and Dring, 1975; Motomura and Sakai, 1981), which was checked under the microscope regularly. To eliminate the confounding factor of red irradiance in our transcriptomic analysis, red light was replaced by white light with a photon fluence rate of 10 - 15 µmol photons m^−2^ s ^−1^ (16hL:8hD) provided by fluorescent lamps OSRAM L 18W for one week prior to and during the experiment. These conditions retained our gametophytes’ cultures at a vegetative stage. At the start of the experiment 3.10 +/−0.10 g of fresh weight of female gametophyte clumps and 2.20 +/−0.10g of male gametophytes clumps were each transferred separately to plastic petri dishes (diameter 9.5 cm × height 3 cm) (n=4) and exposed to three temperatures (4°C, 12°C and 20°C). Experimental temperatures were chosen to mirror mean minimum (4°C) and maximum temperatures (20°C) at Helgoland (Breitbach *et al.*, 2016) and a control temperature (12°C) that falls within optimal growth range for the species (Bolton and Lüning, 1982; Lee and Brinkhuis, 1988). Experiments were conducted in three environmentally controlled rooms and additionally temperatures were kept constant by circulating water baths (models Haake DC1, VWR 1136D). After 14 days of temperature exposure, gametophytes were sampled for RNA extraction. Samples were frozen in liquid nitrogen and stored at −80°C until further use. RNA extraction was performed within three weeks after sampling.

Maximum photosynthetic quantum yield of PS II (Fv/Fm) was measured at the start and end of the experiment (n=4). Gametophytes were dark-adapted for 10 min prior to the measurements. Afterwards, Fv/Fm was measured with an Imaging PAM (Pulse Amplitude Fluorometer; Walz, Effeltrich, Germany).

Fresh weight (n=4) was measured at the beginning and at the end of the experiment with a laboratory scale 40SM-200A (PRECISA, Switzerland). All data were tested for normality using the Shapiro-Wilk normality test and for homogeneity of variances using the Levene’s test. Differences in growth and Fv/Fm were tested using a two-way ANOVA with the fixed factors temperature and sex. Significant differences and interaction of means were compared with the post hoc Tukey test (HSD). All statistical analyses were carried out using SPSS software version 25 (IBM, Armonk, USA). The significance level for all analyses was set at α= 0.05.

### RNA extraction, Illumina sequencing and data processing

Total RNA extraction was conducted using the method described in Heinrich *et al.* (2012). RNA quality was analysed by the NanoDrop ND-1000 UV-Vis Spectrophotometer and Agilent 2100 Bioanalyzer (Agilent Technologies, Germany). Three biological replicates per treatment were sequenced. cDNA libraries were prepared with an Illumina TruSeq RNA Library Prep Kit according to the manufacturer protocol. The libraries were sequenced on an Illumina Hiseq 2500 and 75 bp paired reads were clipped using default values of the Illumina software. Raw reads were quality controlled by FastQC (https://www.bioinformatics.babraham.ac.uk/projects/fastqc/) and quality filtered using Trimmomatic v. 0.36 (Bolger *et al.*, 2014). Quality filtering was performed using the following parameters: leading 3, trailing 3, sliding window 4:15, minlen 30. The Illumina sequence reads generated during the present study are available in the Array express repository under the accession number E-MTAB-8267 (https://www.ebi.ac.uk/arrayexpress/experiments/E-MTAB-8267/).

Reads from all treatments (three temperatures × two sexes) were assembled *de novo* the RNAseq assembler Trinity v 2.4.0 with default parameters including the reads normalization step corresponding to the Trinity implementation of the diginorm method (Grabherr *et al.*, 2011). The quality of the transcriptome assemblies was evaluated by using BUSCO v3.0.2 (Waterhouse *et al.*, 2017) applying an eukaryote dataset (OrthoDB v9.1). Gametophytes reads were pseudo-aligned with Salmon (Patro *et al.*, 2017) against the *de novo* assembled transcriptome, the previously assembled sporophyte transcriptome (Li *et al.*, 2019, Array express:E-MTAB-7348), the available *S. japonica* genome (Ye *et al.*, 2015) and previously assembled transcriptomes of *S. latissima* (Heinrich *et al.*, 2012; Jackson *et al.*, 2017). A PCA plot of the counts-per-million, followed by a log2 transformation, of all treatments was generated by a Trinity script. Differential expression was calculated using DESeq2 (Love *et al.*, 2014) at Trinity’s gene level with an adjusted level of P≤0.001 and a log_2_ fold change of at least 2 indicating significance. Tools were executed using the scripts included in the Trinity package v 2.4.0 (Grabherr *et al.*, 2011). Functional annotation was performed using the Trinotate functional annotation pipeline (Bryant *et al.*, 2017) with the UniRef90 database as additional reference (all databases up to date in October 2017). In addition, annotation of transcripts via DIAMOND v0.9.13 similarity search (Buchfink *et al.*, 2015), using “more-sensitive” option against the NR database (October 2017) was performed.

To investigate the function of significantly up- and down-regulated genes, Gene Ontology (GO) enrichments were conducted using GOseq (Young *et al.*, 2010). Results were summarized with CateGOrizer by applying GO slim (Hu *et al.*, 2008). Venn diagrams were produced applying a webtool (http://bioinformatics.psb.ugent.be/webtools/Venn/).

## Results

### Photosynthetic efficiency and growth

To estimate overall fitness of the gametophytes, we assessed maximum photosynthetic quantum yield by measuring Fv/Fm and fresh weight at the beginning and end of the experiment. There was no significant effect of temperature and sex on fresh weight of gametophytes, measured as percentage of initial at the end of the 14 days exposure (temperature, p=0.159, sex, p=0.525). In all treatments a modest increase in fresh weight up to a maximum of 20% in females at 12°C was observed. No significant treatment changes were detected for Fv/Fm (data not shown).

### *De novo* transcriptome

To explore patterns of gene expression, we sequenced RNA obtained from male and female gametophyte samples exposed to three different temperatures (4°C, 12°C and 20°C). The number of reads per library ranged from 20 to 42 million with an average of 33 million reads. The remapping rate of gametophytes cDNA libraries on the *de novo* assembled transcriptome was on average 86% which is 5 to 18 % better than on other available transcriptomes assemblies (data not shown). The transcriptome assembly consisted of transcripts with more than 1 transcript per million (TPM) in a total of 256,118 transcripts (corresponding to 211,947 Trinity genes) with an average contig size of 713 bp and N50 of 1350 bp. This high total number of genes is a common feature of the Trinity pipeline (e.g. Bryant *et al.*, 2017). Ninety per cent of total expression was present in 58,741 of transcripts with a N50 of 1895 bp. Results of BUSCO analysis indicate that our transcriptome has near-complete gene sequence information for 93% of the genes, 34% complete and single-copy BUSCOs, 59% completed and duplicated BUSCOs, 5% fragmented BUSCOs and 1% missing BUSCOs (Supplementary Table S1). 36,379 transcripts were functionally annotated using the UniProt Swiss-Prot database, 33,606 using Pfam and 60,961 with Uniref90 and 45,240 using NR. A total of 14,240 GO terms and 17,562 KEGG orthology (KO) were assigned to the transcripts.

### Sex-biased gene expression at the control temperature

Several analyses performed in our dataset revealed distinct transcriptomic profiles between male and female gametophytes of *S. latissima*. Gender explained 34% of the variability in the transcriptome as shown in the Principal Component Analysis (PCA) (Figure 1). The second axis of the PCA accounted for 16% of the variability and divided between the three temperatures tested. Many genes were differentially expressed between males and females at the three temperatures (23,698 – 11.2%; Figure 2). Differential gene expression between sexes at the control temperature (12°C – 13,991 DEGs) was higher than at a temperature of 20°C (10,590) and at 4°C (9,411; Figure 2). Differential expression between sexes was more similar between 4°C and 12°C than between 12°C and 20°C or 4°C and 20°C (Figure 3). Across temperatures, a female-biased gene expression was observed, with more up-regulated DEGs in females compared to males. Among DEGs up-regulated in females, 12% were up-regulated at the three temperatures, 29% were uniquely up-regulated at 20°C, 28% at 12°C and 9% at 4°C. In turn, among DEGs up-regulated in males, only 7% were consistent across temperatures, 29% were specifically regulated at 4°C, 28% at 12°C and 29% at 20°C (Figure 3). Consistently, several GO terms were significantly enriched in the comparison between gender and patterns shifted with temperature (Figures 4 to 7). The lower temperature, 4 °C drove more metabolic reorganization in males than females and the opposite held true at 12°C and 20°C for females. Taken together, these results reveal a clear sex-biased gene expression that was modulated by temperature.

**Figure 1.**
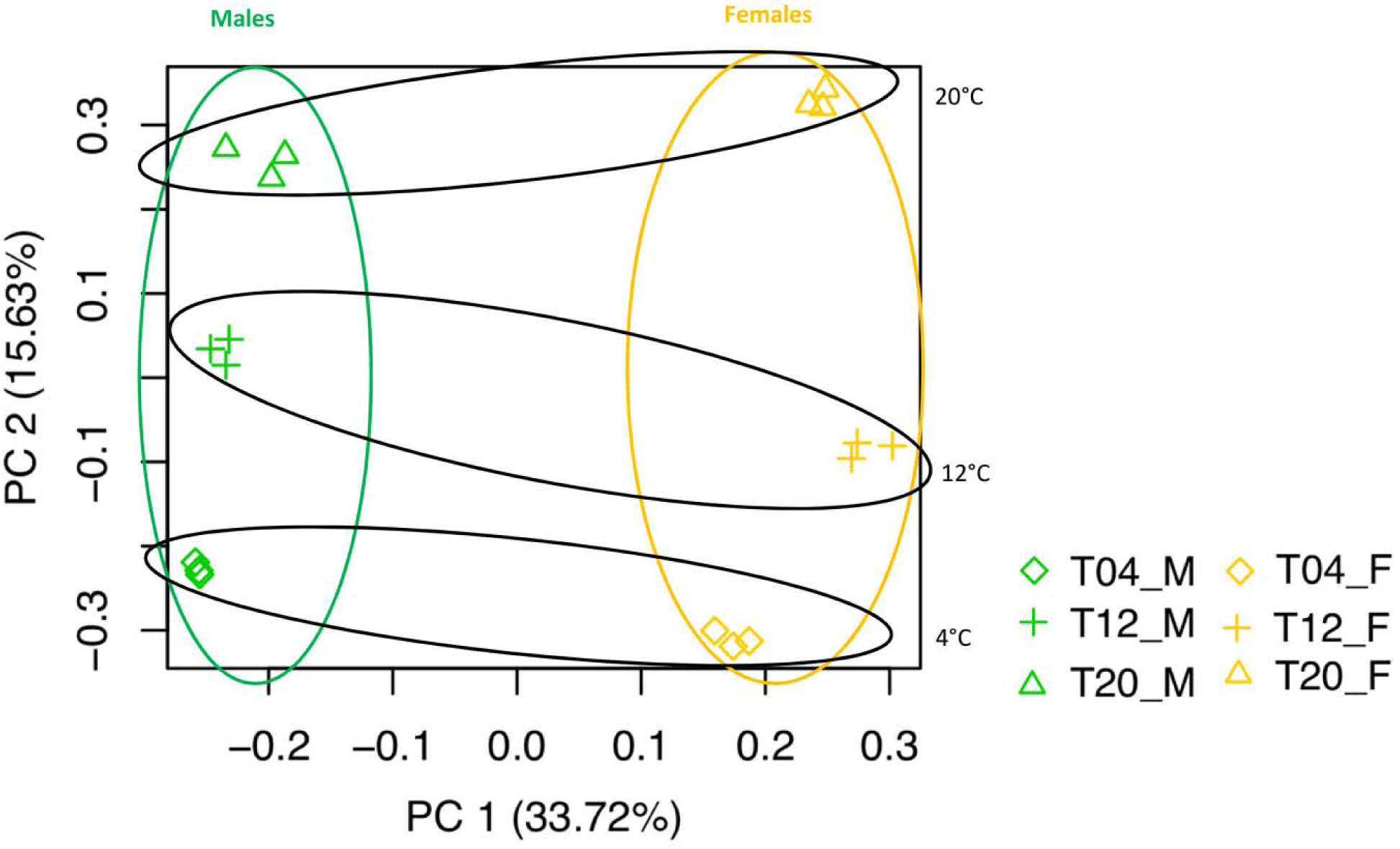
Principal Component Analysis of the counts per million in the treatments (n=3) after 14 days in the experimental temperature treatment. Orange: female samples, green: male samples

**Figure 2.**
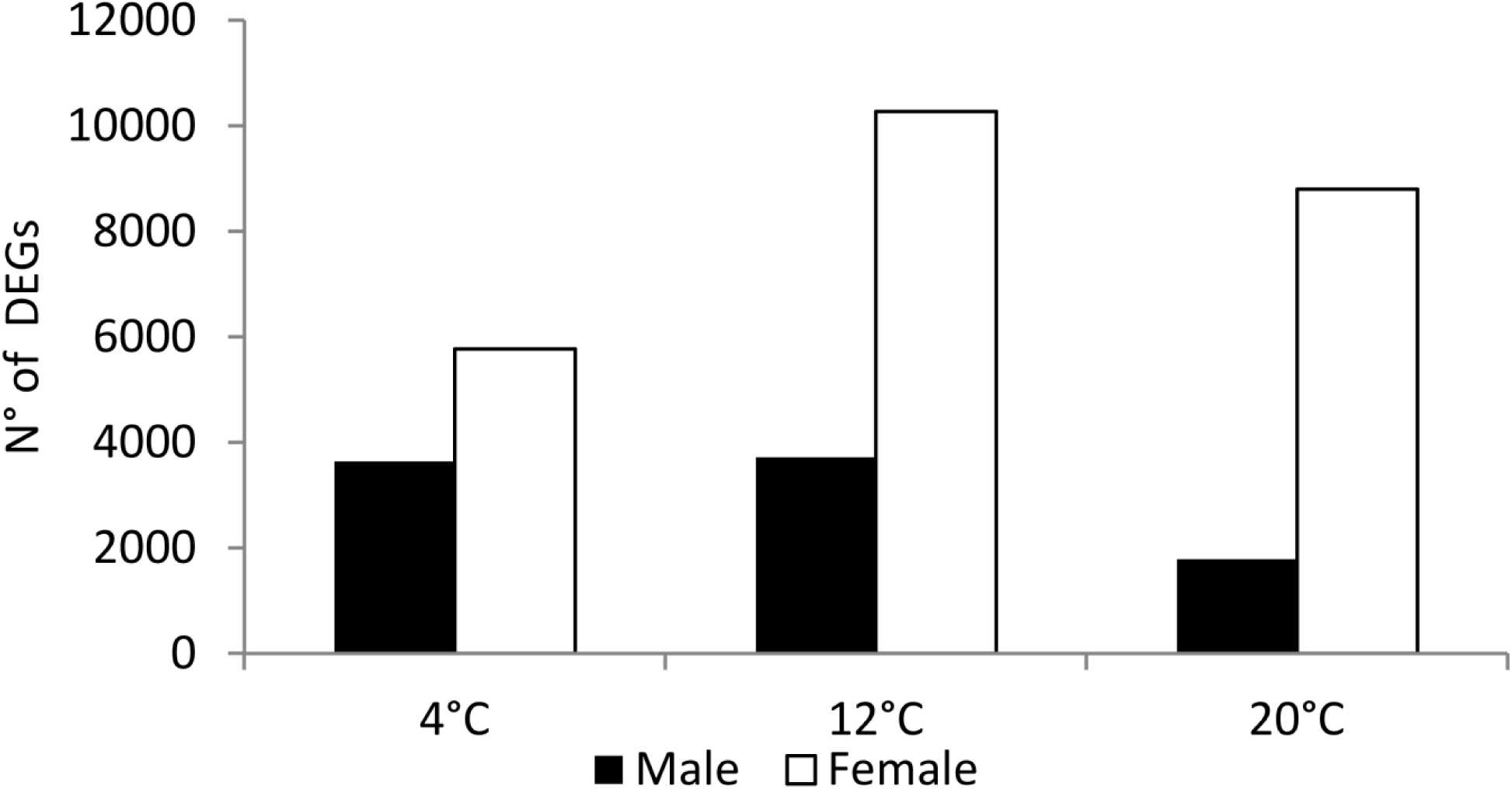
Total number of significant Differentially Expressed Genes (DEGs) (Deseq2, *p* < 0.01; FC > 2) in *Saccharina latissima* gametophytes. Up-regulated DEGs in male compared to female gametophytes are indicated in black, and white bars, respectively.

**Figure 3.**
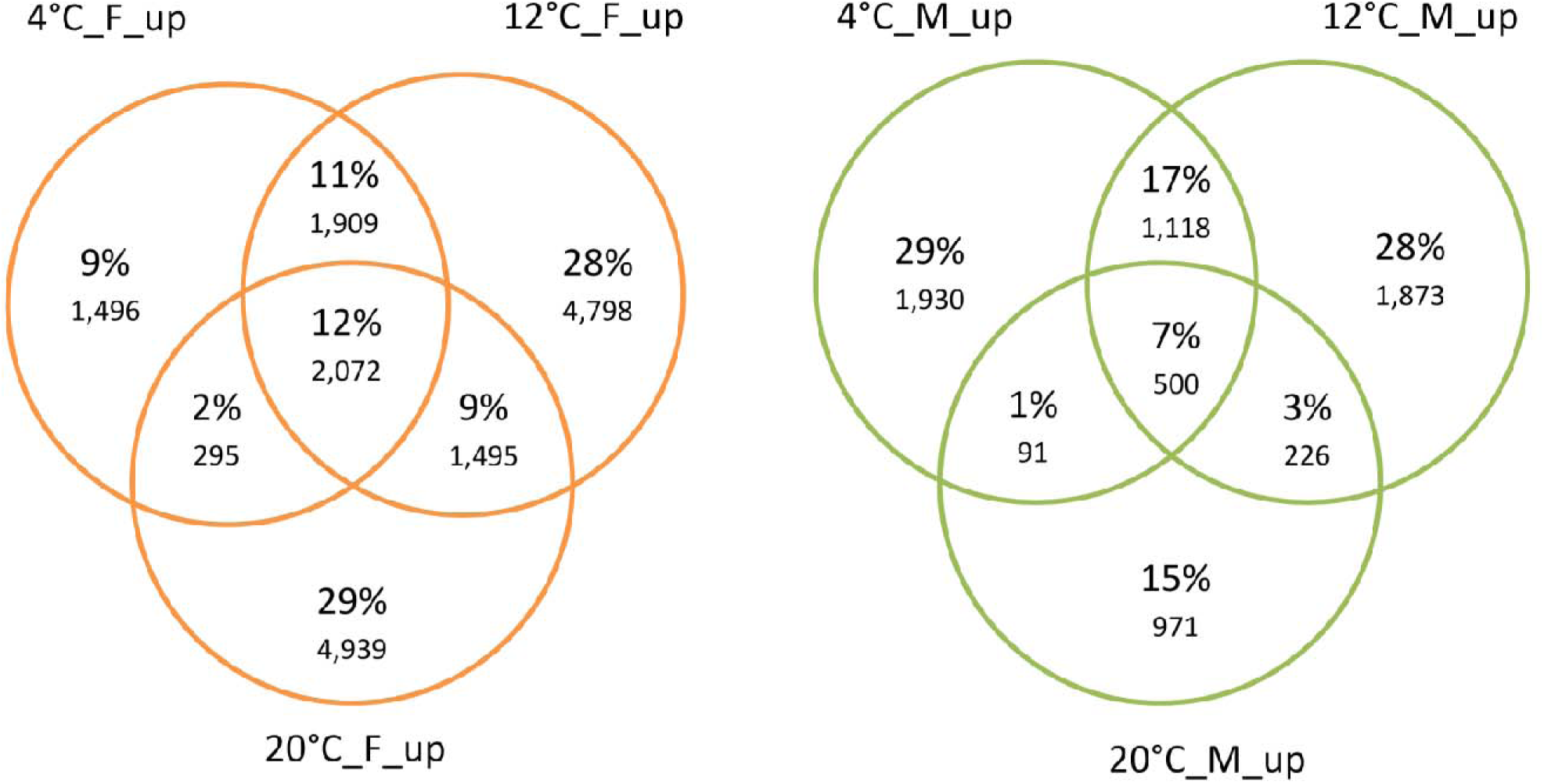
Venn diagrams of DEGs between sexes of *Saccharina latissima* by temperature (p < 0.001; log2FC > 2). Left and orange: DEGs up-regulated in females, Right and green: DEGs up-regulated in males by temperature.

**Figure 4.**
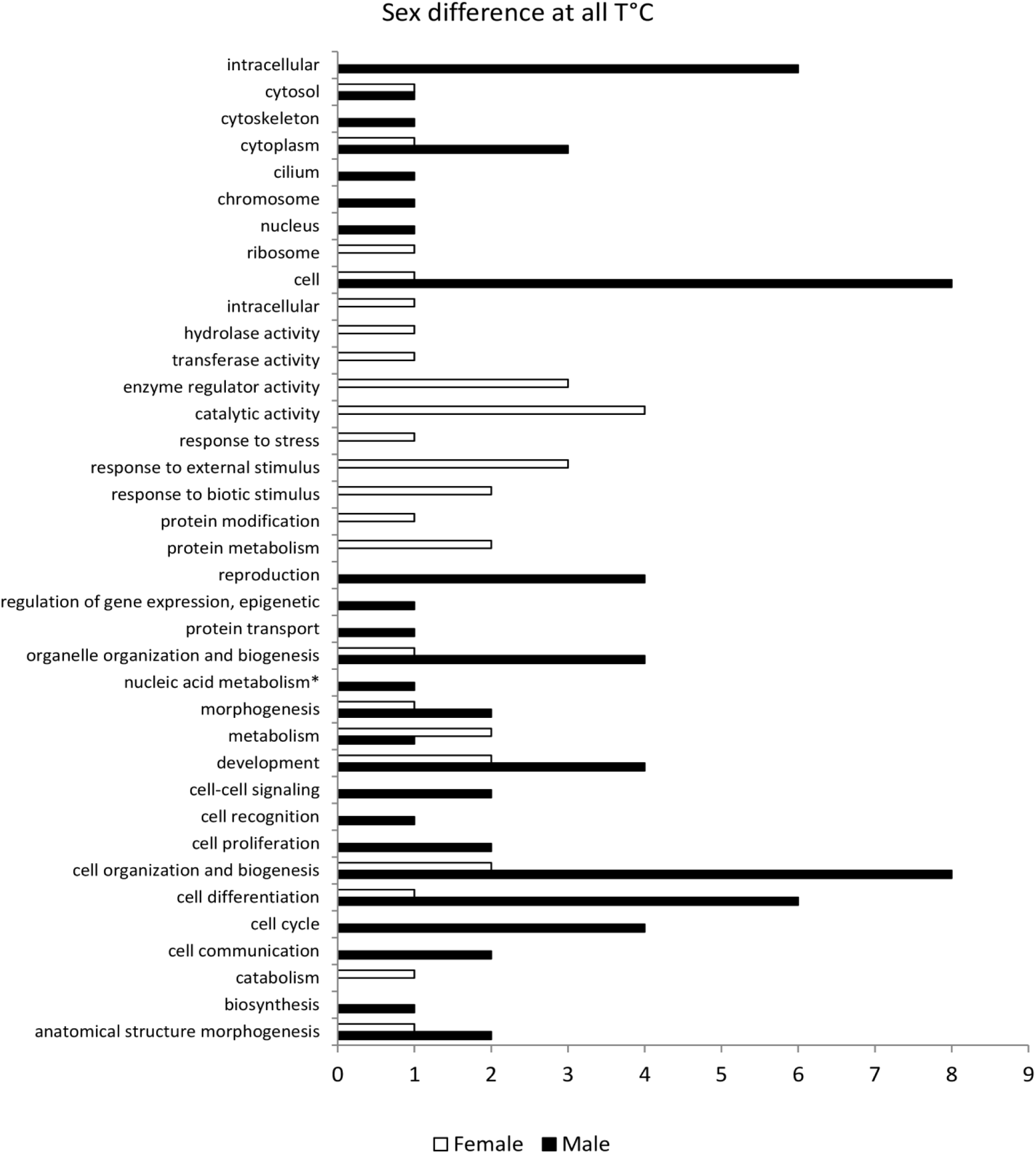
Over-represented GO terms of DEGs consistently up-regulated in female (white) and male (black) gametophytes of *Saccharina latissima* across all temperatures, according to the categories of GO slim. Nucleic acid metabolism* stands for nucleobase, nucleoside, nucleotide and nucleic acid metabolism.

### Putative sex marker genes

Analysis of the genes with the highest contribution to the Principal Component axis 1, measured by its loading values (Legendre and Legendre, 2012), revealed genes potentially related to sex determination in *S. latissima* (Supplementary Table S2). Namely, one gene highly expressed in females and another one highly expressed in males both match one gene coding for homoaconitate hydratase that has been described as female-specific in *Ectocarpus spp.* and *S. japonica* (Lipinska *et al.*, 2017). Also, a gene identified in six other brown algal species as male-specific, the high mobility group protein 1 (Lipinska *et al.*, 2017), was also male-specific in our data. Moreover, we identified two protein kinases: dual specificity protein kinase TTK and serine/threonine-protein kinase OSR1 associated with male gene expression. Two other genes functioning in protein modification, ubiquitin-conjugating enzyme E2 D2 and galactose-3-O-sulfotransferase 3, were correlated with male samples. A gene coding for serine protease/ABC transporter B family protein tagC and a gene coding for calcium channel YVC1 were associated with females.

### Functional annotation of sex-biased genes

To identify the functions of sex-biased genes, we performed a GO enrichment analysis in the set of genes consistently up-regulated in females (2,072) and males (500) across temperatures (Figure 4) and on the full set of genes differentially expressed between sexes at 4°C, 12°C and 20°C (Figures 5-7). Sex-biased genes identified in this study were associated with several metabolic processes. Results of the GO enrichment analysis revealed that at 12°C and 4°C the top 15 overrepresented GO terms were related to general metabolism, such as “peptide biosynthetic process”, “organonitrogen compound biosynthetic process” and “translation” for both females and males up-regulated DEGs (Supplementary Table S3). Terms related to cytoskeleton and signalling, e.g. “cellular component assembly”, “actin filament organization”, “small GTPase mediated signal transduction” were uniquely overrepresented in males while GO terms related to aminoacid metabolism (“carboxylic acid metabolic process”) and nucleic acid metabolism (“nucleoside triphosphate metabolic process”) were uniquely overrepresented in females at both 4°C and 12°C. Considering the extensive list of enriched GO terms and that it contained several human-related biological processes, we applied GO slim to provide an overview. Among GO slim functional categories exclusively over-represented in females consistently across temperatures, we identified GO terms related to metabolism and responses to cues. In the category metabolism, the GO terms “protein metabolism” and “catabolism” were over-represented, while response to cues included the GO terms “response to stress”, “response to external stimulus” and “response to biotic stimulus” (Figure 4). Accordingly, we identified considerably more stress response related DEGs up-regulated in females at the three different temperatures, namely heat-shock and antioxidant proteins (Supplementary Table S4). In turn, functional categories solely over-represented in males included “cell-cell signalling”, “cell cycle” and “cell proliferation”. Moreover, some categories were over-represented in both males and females and could be connected to processes that we expect to take place in both sexes during gametophyte growth and development, namely “cell differentiation” and “cell organization and biogenesis”.

**Figure 5.**
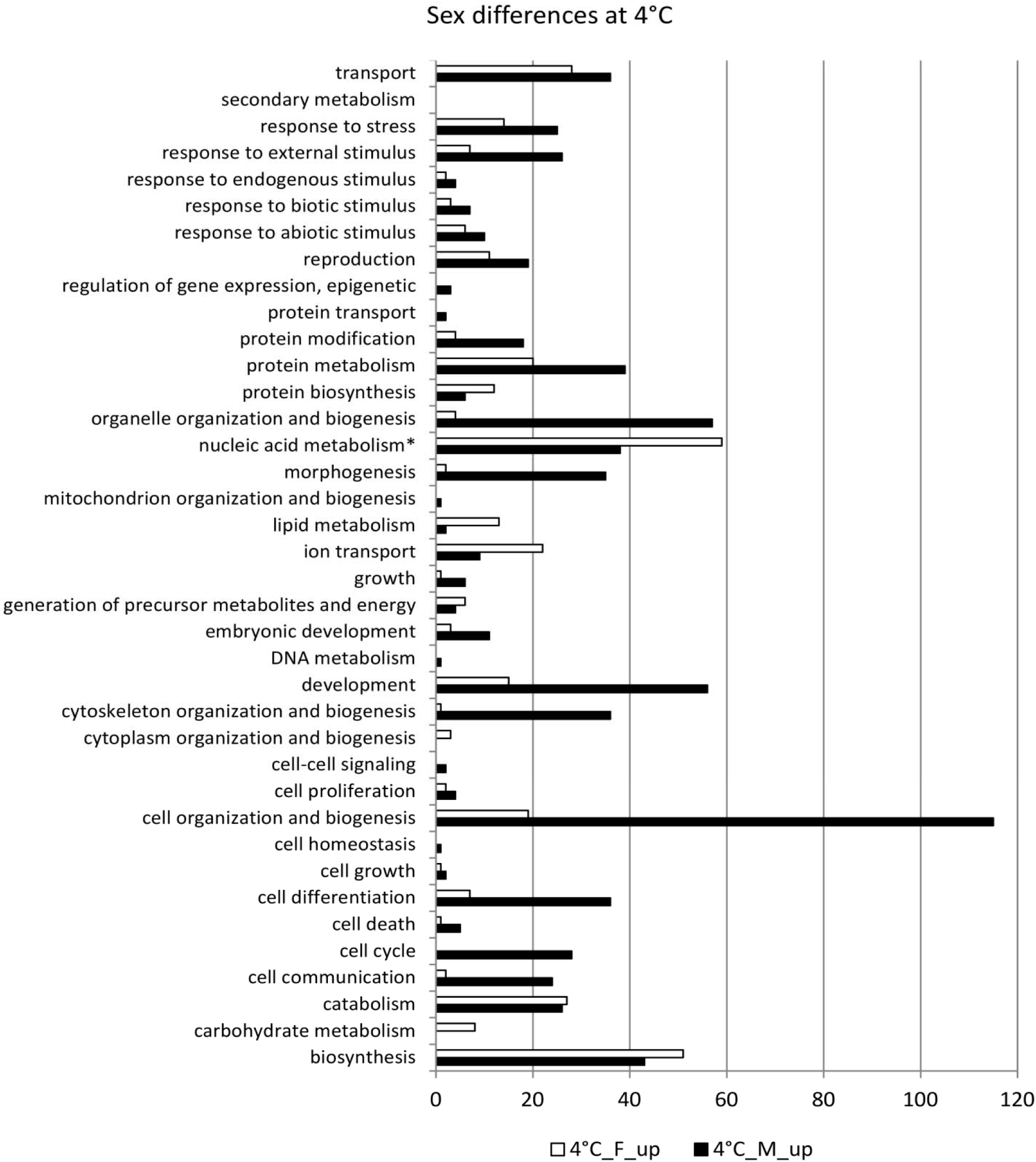
Over-represented GO terms belonging to the ontology Biological Process in male (black) and over-represented in female (white) gametophytes of *Saccharina latissima* at 4°C according to the categories of GO slim. Nucleic acid metabolism* stands for nucleobase, nucleoside, nucleotide and nucleic acid metabolism. Total for the category metabolism - Females: 170, Males: 153.

### Sex-biased gene expression at low and high temperatures

Shifts in the expressed metabolic pathways were observed in response to changes in temperature. The low temperature tested, 4°C, drove more metabolic reorganization in males than in females (Figure 5). “morphogenesis”, “reproduction”, “transport” and responses to endogenous, biotic and abiotic stimulus gathered more enriched GO terms in males than in females at 4°C (Figure 5), and the reverse was observed at 12°C (Figure 6). However, biological processes such as lipid and carbohydrate metabolism and “ion transport” were proportionally higher in females at 4°C and at 12°C than males. Concomitantly, “cell differentiation”, cell and organelle organization and biogenesis were consistently over-represented in males compared to females at both 4°C and 12°C.

**Figure 6.**
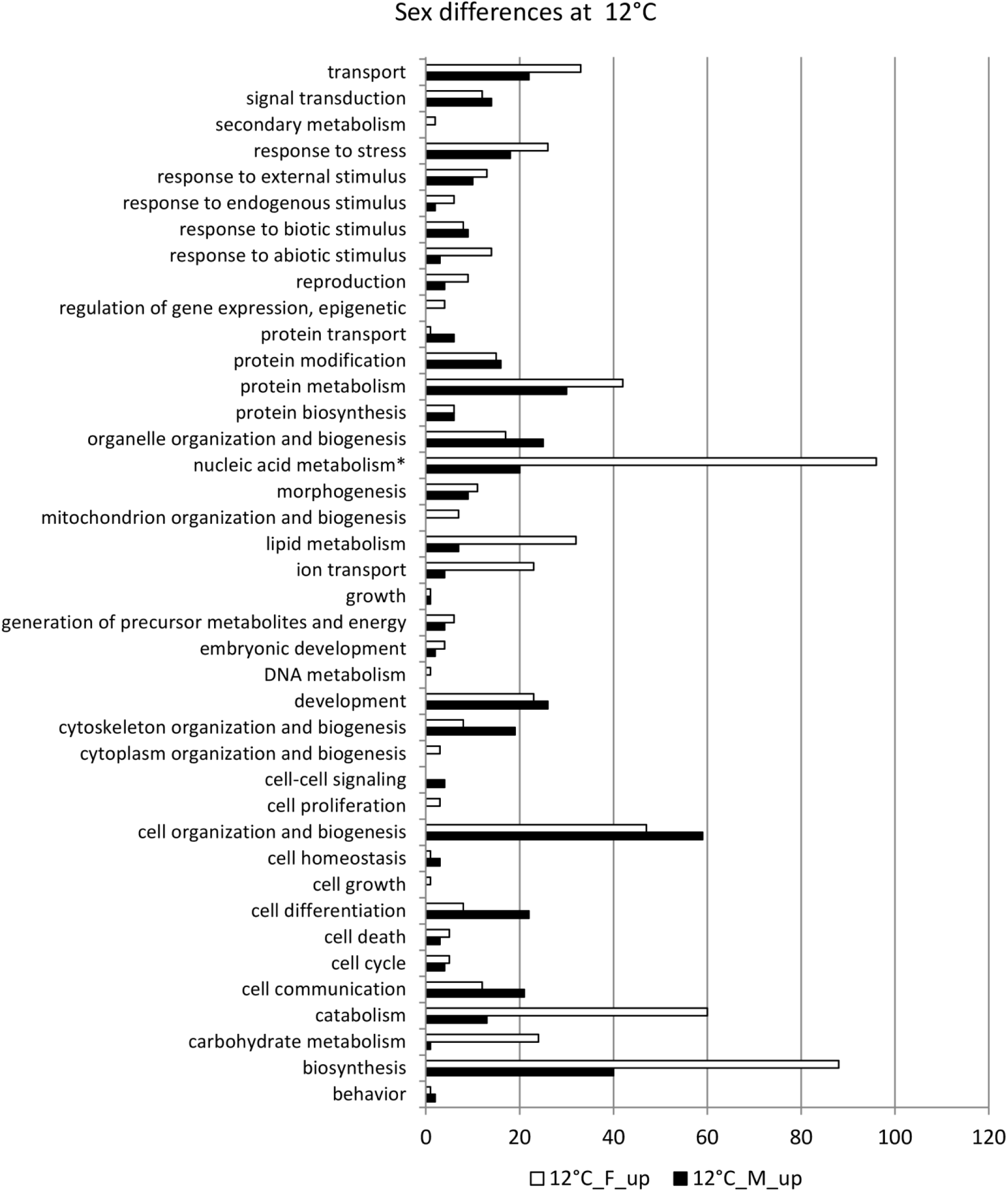
Over-represented GO terms belonging to the ontology Biological Process in male (black) and over-represented in female (white) gametophytes of *Saccharina latissima* at 12°C according to the categories of GO slim. Nucleic acid metabolism* stands for nucleobase, nucleoside, nucleotide and nucleic acid metabolism. Total for the category metabolism - Females: 354, Males: 120.

At 20°C, we observed a decrease in the differentiation between sexes, revealed by the lower number of enriched GO terms within the root biological processes differentially expressed between males and females (Figure 7). Almost all the functional categories were over-represented in females: “transport”, “signal transduction”, “response to stress”, “protein metabolism”, “cell organization and biogenesis” and “cell communication” gathered the higher number of GO terms. The top 15 over-represented GO terms in females included “superoxide anion generation”, “positive regulation of oxidative stress-induced intrinsic apoptotic signalling pathway”, “transposition”, “DNA integration” (Supplementary Table S3).

**Figure 7.**
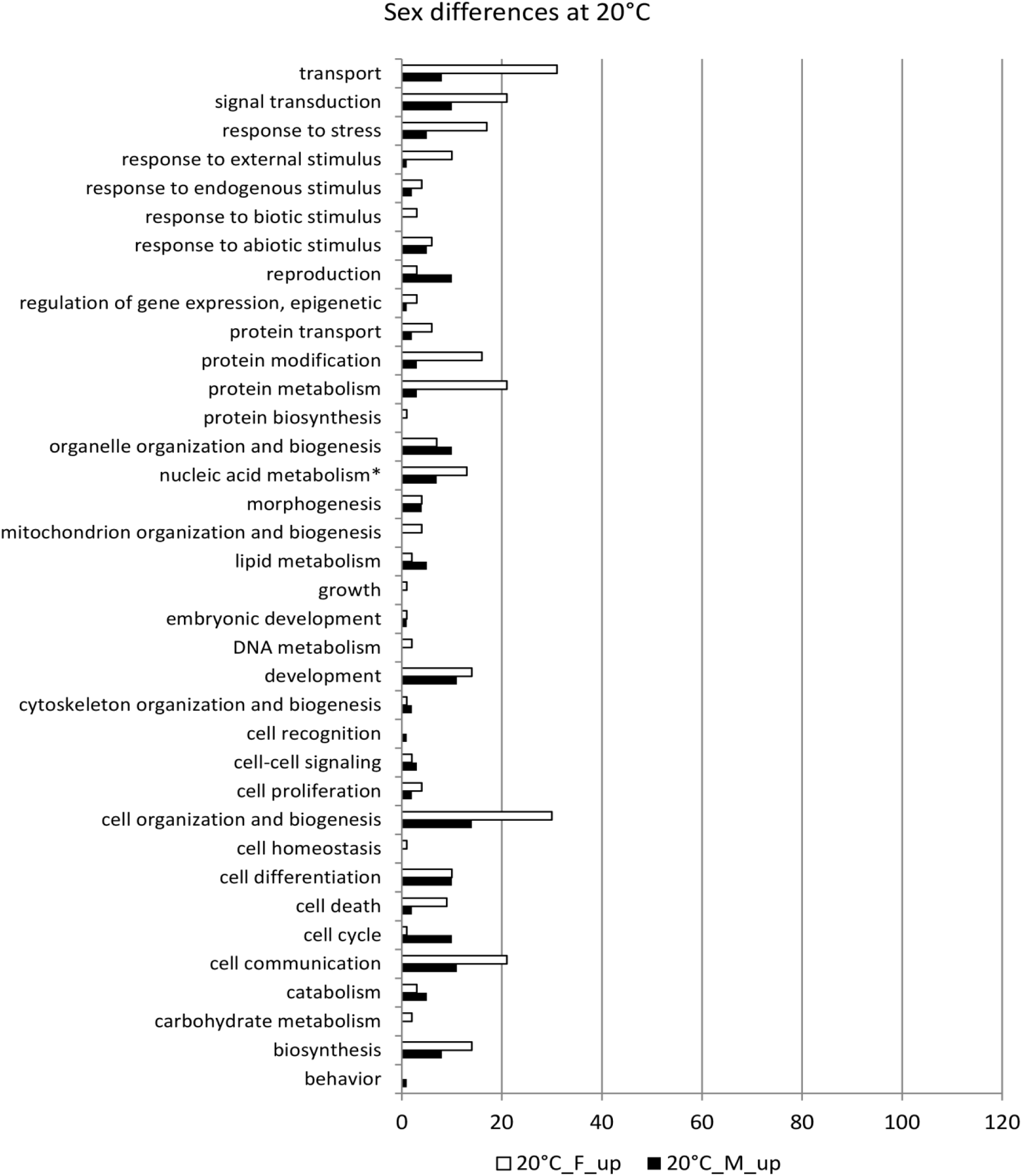
Over-represented GO terms belonging to the ontology Biological Process in male (black) and over-represented in female (white) gametophytes of *Saccharina latissima* at 20°C according to the categories of GO slim. Nucleic acid metabolism* stands for nucleobase, nucleoside, nucleotide and nucleic acid metabolism. Total for the category metabolism - Females: 55, Males: 17

## Discussion

### Sex-biased gene expression underlies gender-specific metabolic needs

In this study, we identified a high number of genes differentially expressed between male and female gametophytes which were in the vegetative stage. This pattern is consistent with descriptions for *Ectocarpus* sp. by Lipinska *et al.* (2015) who reported that differential gene expression between males and females was higher during the immature stage than during the fertile one. Furthermore, more female-than male-biased genes were identified, which agrees with results obtained during gametogenesis in *S. latissima* (GA Pearson *et al*. 2019, unpublished data). However, this is in contrast to the situation in two other brown algae, *Fucus vesiculosus* (order Fucales) (Martins *et al.*, 2013), and *Ectocarpus* sp. (order Ectocarpales) (Lipinska *et al.*, 2015). Although there are contrasting patterns in sex-biased gene expression within brown algae, the three species belong to different orders and even high variability in sex-biased gene expression among species of the same genus has been described before (Metta *et al.*, 2006). Moreover, differences may mirror the different life cycles (Coelho *et al.*, 2019). While Laminariales and Ectocarpales have a haplo-diplontic life cycle, Fucales display a diplontic life cycle. Moreover, the degree of sexual dimorphism in the haplo-phase (gametophytes) of Laminariales is stronger than in the order Ectocarpales and while the first produce anisogamous gametes with a large non-motile female egg cell and small motile male sperms, Ectocarpales gametes are both motile and isogamous (Luthringer *et al.*, 2014).

Functional analysis of the differentially expressed genes revealed that several metabolic pathways are distinctly regulated between female and male gametophytes. In females, several GO terms related to general metabolism are enriched, such as “carbohydrate metabolism”, “lipid metabolism”, “nucleobase, nucleoside, nucleotide and acid metabolism”. In addition, the over-represented GO term “generation of precursor metabolites and energy” suggest that female gametophyte cells are actively investing in cell growth and energy production. This is consistent with observational microscopy studies, as female gametophyte cells tend to grow in size while male gametophyte cells tend to grow in number (Destombe and Oppliger, 2011; Lüning, 1980; Park *et al.*, 2017). This investment in cell growth might be advantageous for later sporophyte growth as larger female gametophyte cells will release bigger eggs that tend to generate larger sporophytes (Bell, 1997). Within lipid metabolism, we identified four genes coding for lipoxygenase that were sex-specific in *S. latissima* in this study (Supplementary Table S5) and during gametogenesis (GA Pearson *et al*. 2019, unpublished data), as in *Ectocarpus* sp. (Lipinska *et al.*, 2015) although female-specific in the first and male-specific in the latter species.

Lipoxygenases are involved in lipid oxidation and play an important role in the biosynthesis of oxylipins - lipophilic mediators that have several physiological functions such as senescence, growth and development, tolerance to stress and cell homeostasis (Hou *et al.*, 2015; Maynard *et al.*, 2018). In brown algae, they are involved in the defense response to pathogens and grazing (Cosse *et al.*, 2009; Ritter *et al.*, 2017) and their synthesis was induced under copper stress (Ritter *et al.*, 2008). Considering the sex-biased gene expression of lipoxygenases in both *Ectocarpus* sp. and *S. latissima* we hypothesize that they are involved in regulating sexual differentiation in brown algae. Further work is required to understand the specific function of lipoxygenases in sexual differentiation and its different expression between brown algae species. In addition, two genes coding for mannuronan C5-epimerase were female-specific and one was male-specific in *S. latissima*, while in *Ectocarpus* sp. two genes were male and one was female-specific. Mannuronan C5-epimerases belong to large multigenic families that are involved in the synthesis of alginate, a major component of cell walls in brown algae (Michel *et al.*, 2010). Previously, Fischl *et al.* (2016) reported four mannuronan C5-epimerases genes preferentially expressed in Ectocarpales gametophytes over sporophytes, suggesting a role in the gametophyte development. Differential expression of some of the genes enconding this enzyme might contribute to the different morphologies of the male and female gametophytes’ cells.

A closer inspection of the functional annotation of DEGs revealed that some genes involved in oxidative stress responses were up-regulated in female gametophytes of *S. latissima*. Similar patterns were also reported for *S. latissima* during gametogenesis (GA Pearson *et al*. 2019, unpublished data) and in the female gametophyte of *Arabidopsis* (Wuest *et al.*, 2010). Reactive oxygen species play a role in embryo sac development and fertilization in *Arabidopsis* (Martín *et al.*, 2013), cell division (Livanos *et al.*, 2012a) and cytoskeleton homeostasis (Livanos *et al.*, 2012b) in plants. Thus, we hypothesize that reactive oxygen species might function as signaling molecules regulating gametophyte development, as previously reported in gender-biased gene expression in primates and mice (Blekhman *et al.*, 2010; Yang *et al.*, 2006), suggesting that these are highly conserved processes.

In vegetative males, GO terms in the GO slim functional categories “cell communication” and “cell-cell signaling” were over-represented which is probably connected to the fact that gametophyte maturation is dependent on both environmental factors and biotic cues, namely the release of pheromones (Lüning and Müller, 1978). Signal transduction might be crucial to allow for temporal and spatial overlap in gametophyte maturation which is critical for successful fertilization in kelps (Schiel and Foster, 2006). During our experiments no reproductive structures were visible and males did not receive female cues for maturation as sexes were kept separated. Nevertheless, our data suggests that male gametophytes prepare in advance for maturation at the transcriptomic level probably to be able to readily synchronize gamete release with females once cues are received. This might confer a competitive advantage and ensure successful fertilization. Li *et al.* (2013) have observed that the best fertilization rates in *S. japonica* occur 60-120 minutes after male gamete release and their viability is limited to 12 hours. This is further supported by the expression of a “flagellar associated” gene reported from *Ectocarpus* which was only expressed in males of *S. latissima* at 4°C and 12°C. Two “flagellar associated proteins” also displayed male-biased gene expression in immature gametophytes of *Ectocarpus* (Lipinska *et al.*, 2015).

Furthermore, higher enriched GO terms within the categories “cell”, “cytoskeleton” and “organelle organization and biogenesis” are in accordance with cell division being enhanced in male gametophyte filaments and these being more highly branched than female gametophyte filaments. Among these functional categories, we found two genes encoding DNA helicases with very high expression in males. DNA helicases are evolutionary conserved among eukaryotes and involved in DNA replication, namely by positive regulation of telomere length, replication of highly transcribed RNA polymerase II genes and unwinding G□quadruplex DNA structures (Rhodes and Lipps, 2015; Sabouri, 2017). Therefore, higher expression of these genes in males might be connected to higher cell division rates.

### Changes in temperature modulated sex-biased gene expression

Functional analysis revealed that sex-biased gene expression shifted with temperature. Differences between sexes were less pronounced at 20°C than at 4°C and 12°C suggesting that the gametophytes alter their metabolism from developmental and sex differentiation processes to a heat stress protection response that involves similar pathways in both sexes. This is in agreement with 4°C and 12°C being more suitable temperatures for gametophyte growth and gametogenesis than 20°C (Lüning, 1980).

It is widely accepted that temperature is a major factor controlling reproduction in seaweeds (Andrews *et al.*, 2014; Egan *et al.*, 1989; Hurd *et al.*, 2014a). This is further supported in our study by the interactive effect of temperature and sex in gene expression patterns in *S. latissima*. Only a small proportion of differentially expressed genes were consistent across temperatures (12% for females and 7% for males) and females responded stronger to the higher temperature (20°C) while more genes were overexpressed in males at lower temperatures (4°C). The effect of temperature on growth of *S. latissima* gametophytes has been previously demonstrated: Lüning (1980) observed optimum vegetative growth between 10 and 19°C and gametophyte death after one week at 22°C. Another study revealed that gametophytes survived 22°C and even 23°C but with visible cell damage (Bolton and Lüning, 1982). Hence, the higher temperature tested in the present study (20°C) is very close to the upper survival temperature in gametophytes and to the maximum temperature reached during summer at the original collection site of the gametophytes (Helgoland, North Sea, (Breitbach *et al.*, 2016)). Other studies have suggested that male kelp gametophytes are slightly more thermotolerant than females (Lee and Brinkhuis, 1988; tom Dieck, 1993). This is supported by the differential gene expression of our study. The male gametophytes seem to be more resilient to the heat stress as functional categories such as “response to stress”, response to external and abiotic stimuli and “cell death” were over-represented in females and categories such as “cell cycle” were over-represented in males. In addition, induction of transposable elements by stress and their effects on gene regulation has been observed in plants, namely in response to heat stress (Dubin *et al.*, 2018; Makarevitch *et al.*, 2015). The over-representation of GO terms related to transposition in females at 20°C suggests a role in heat stress response in *S. latissima* gametophytes. Concomitantly, other enriched GO terms connected to stress response, such as “superoxide anion generation”, “positive regulation of oxidative stress-induced intrinsic apoptotic signalling pathway” were also among the top 15 over-represented GO terms in females at 20°C. This further supports the higher sensitivity to heat stress of female gametophytes. Assessing differential sensitivity to high temperatures between sexes in gametophytes is relevant for hybridization protocols aiming to produce more tolerant cultivars in aquaculture, as it is already current practice in crop plant species (Zinn *et al.*, 2010). Recently, Martins *et al.* (2019) demonstrated that for the kelps *Laminaria digitata and L. pallida* female parent gametophytes contributed more for the thermal response of the hybrids than males. Our work suggests that the same might be true for *S. latissima* and informs current ongoing efforts to maximize sugar kelp cultivation practices.

Shifts in sex ratio in response to sub-optimal temperatures have already been described in several kelp species (Izquierdo *et al.*, 2002; Nelson, 2005; Oppliger *et al.*, 2011). In *S. latissima*, Lee and Brinkhuis (1988) described a greater proportion of male compared to female gametophytes at higher temperatures (between 17°C and 20°C). Female gametophyte grew at 20°C, but with a lower rate than between 4 and 17°C and their fecundity was repressed. Pronounced temperature-mediated sex ratio shifts might lead to population declines in several species, especially at sites where species are close to their upper thermal tolerance limit, and may have an impact on the range distribution of species (Hays Graeme *et al.*, 2017; Janzen, 1994; Ospina-Alvarez and Piferrer, 2008). Differences in the sex ratio between populations along the distributional range have been observed for several plants (Barrett and Hough, 2012) and for the kelp *Lessonia nigrescens* (Oppliger *et al.*, 2011), which warrants concern that skewed sex ratios due to global warming might compromise populations viability. At the gametophyte stage, global warming may lead to a mismatch between male and female gametophyte performance that might compromise sexual maturation and fertilization, as has been recently highlighted by de Bettignies *et al.* (2018). At the same time, the high levels of gene expression reported here indicate that vegetative gametophytes are still metabolically active rather than “dormant” as in plant seeds (Schiel and Foster, 2006) and therefore more vulnerable to abiotic stress. Therefore, the sex differences in gene expression patterns in response to temperature described here for *S. latissima* prompts for further investigation on the vulnerability of the gametophyte life-stage to temperature and its potential effects in reproductive success and phenology.

## Supplementary data

Supplementary Table S1: Results of BUSCO analysis of transcriptome completeness

Supplementary Table S2: Top 100 genes contributing to PC1 of the Principal Component analysis according to their loading values

Supplementary Table S3: Top 15 over-represented GO terms by sex (male, female) and temperature (4°C, 12°C and 20°C), both total GO terms and sex-exclusive.

Supplementary Table S4: List of stress response related DEGs between sexes at the three different temperatures

Supplementary Table S5: List of DEGs between sexes found in common in *Ectocarpus sp.* And *S. latissima*

## Acknowledgments

This work was supported by the German Research Foundation for funding within the ERA-Net Cofund BiodivERsA 3 program MARFOR (ANR-16-EBI3-0005-01). Further funding was provided by the MARES Joint Doctoral Programme on Marine Ecosystem Health & Conservation funded through Erasmus Mundus (grant number MARES_14_09) and by the Alfred-Wegener-Institute Helmholtz-Centre for Polar and Marine Research (Bremerhaven, Germany).

